# Reduced cortical VPS26B levels are associated with altered glutamate receptor expression and synaptic protein loss in the primary motor cortex of a Parkinsonian mouse model

**DOI:** 10.64898/2026.05.18.726103

**Authors:** Thanh Thi Hai Nguyen, Jung Bae Seong, Jincheol Seo, Jinyoung Won, Se-Hee Choe, Hae Rim Kim, Ki-Hoan Nam, Young-Hyun Kim, Youngjeon Lee

**Affiliations:** National Primate Research Center, Korea Research Institute of Bioscience and Biotechnology (KRIBB), Republic Of Korea; Department of Functional Genomics, KRIBB School Of Bioscience, Korea University Of Science and Technology, Republic Of Korea; Laboratory Animal Resource Center, Korea Research Institute of Bioscience and Biotechnology (KRIBB), Republic Of Korea

## Abstract

Parkinson’s disease (PD) is associated with motor impairment and cortical synaptic dysfunction, which involve altered glutamate receptor trafficking, yet the underlying mechanisms remain incompletely understood. VPS26B, a component of the retromer complex, regulates GluA1 recycling in the trans-entorhinal cortex region. However, its role in the primary motor cortex (M1) under Parkinsonian conditions has not been explored. Here, we show that VPS26B levels are reduced in the M1 of an MPTP-induced PD mouse model, accompanied by decreased surface GluA1 and synaptic protein levels. VPS26B overexpression partially attenuated these alterations. In the accelerating rotarod test, VPS26B-deficient mice exhibited unstable motor performance following MPTP administration, whereas VPS26B overexpression was associated with improved performance in both wild-type and knockout mice. These findings suggest that cortical VPS26B may contribute to maintaining glutamate receptor surface expression and synaptic protein levels, especially under Parkinsonian conditions, with potential implications for motor learning.

## 1. Introduction

Parkinson’s disease (PD) is classically characterized by dopaminergic degeneration in the substantia nigra (SN) [1]. However, accumulating evidence indicates that PD is also associated with cortical dysfunction, including altered glutamatergic signaling and impaired synaptic plasticity [2, 3]. Consistent with this, emerging evidence suggests that cortical dysfunction may contribute to dopaminergic neuron vulnerability in PD [4].

Synaptic plasticity in the primary motor cortex (M1) plays a critical role in motor learning. Patients with PD exhibit impaired motor learning associated with reduced long-term potentiation (LTP)-like plasticity [5, 6]. Importantly, such plasticity deficits are detected early in PD and may reflect cortical dysfunction contributing to motor impairment [7].

Cortical synaptic plasticity relies on glutamate receptor signaling, particularly AMPA-type ionotropic glutamate receptors (AMPARs), which mediate the majority of fast excitatory synaptic transmission [8, 9]. Among these, GluA1-containing AMPARs are essential for activity-dependent synaptic strengthening via regulated recycling and insertion into the postsynaptic membrane [9, 10]. This process determines whether internalized receptors are recycled to the plasma membrane or targeted for degradation [11]. However, the mechanisms underlying GluA1 recycling and surface maintenance in the M1 remain incompletely understood, particularly in the context of PD.

Retromer is a multimeric protein trafficking complex that transports protein cargo from early endosomes to the trans-Golgi network or the plasma membrane [12, 13]. By directing cargo back to the plasma membrane and preventing its lysosomal degradation, retromer plays a critical role in maintaining protein homeostasis [14, 15]. Disruption of retromer components leads to cargo trapping within endosomal compartments and increased lysosomal degradation, processes that have been implicated in several neurodegenerative diseases [16–19]. In PD, mutations in the core retromer component Vacuolar protein sorting 35 (VPS35) are linked to dopaminergic neurodegeneration [20–23]. However, the roles of other core retromer components and retromer-dependent recycling mechanisms in PD pathophysiology have been less extensively studied.

Vacuolar protein sorting 26B (VPS26B) is a core component of the retromer complex that regulates GluA1 recycling in memory-related cortical regions in the context of Alzheimer’s disease [24]. These findings raise the possibility that VPS26B may regulate AMPAR trafficking beyond memory-related regions. The role of VPS26B in maintaining glutamate receptor expression and synaptic integrity in the M1 under PD-like conditions remains unknown. Here, we hypothesized that VPS26B loss in the M1 is linked to altered GluA1 surface expression and synaptic dysfunction, potentially contributing to impaired motor performance.

In this study, we show that VPS26B levels are reduced in the M1 of an MPTP-induced mouse model of PD. This reduction coincides with decreased surface expression of GluA1 and loss of synaptic proteins involved in cortical synaptic function. AAV-mediated VPS26B overexpression prior to MPTP administration preserves GluA1 surface availability and attenuates synaptic protein loss. At the behavioral level, VPS26B overexpression attenuated deficits in rotarod performance. Together, these findings suggest that VPS26B contributes to cortical molecular alterations under Parkinsonian conditions.

## 2. Materials and methods

### 2.1. Animals

Animals were housed at a constant room temperature (RT) with free access to water and food in a 12h light-dark cycle in the animal facility of Ochang Laboratory Animal Resource Center, Korea Research Institute of Bioscience and Biotechnology, Korea. Bedding was changed weekly until sacrifice. All experimental procedures were approved by the Institutional Animal Care and Use Committee of KRIBB (No. KRIBB-AEC-26051) and conducted in accordance with KRIBB IACUC and ARRIVE guidelines [25]. Mixed-sex animals were used, and sex was not used as an experimental variable in this study. Nine-week-old C57BL/6J wild-type (WT) and VPS26B knockout (KO) mice were randomly assigned into five groups: (G1) WT control, (G2) WT + MPTP, (G3) WT + VPS26B + MPTP, (G4) KO + MPTP, (G5) KO + VPS26B + MPTP (Supplementary Table 1).

The planned sample size was six mice per group (control group: n = 4). Due to the limited availability of KO mice, group G5 consisted of four animals. During the experiment, two animals in G3 and three animals in G4 died following MPTP administration, consistent with previous reports describing the systemic toxicity of the MPTP model [26]. Therefore, the final sample sizes used for analysis were n = 4 (G1), n = 6 (G2), n = 4 (G3), n = 3 (G4), and n = 4 (G5). A saline-treated KO group was not included as the present study was primarily designed to investigate VPS26B-associated alterations in PD-like conditions.

### 2.2. Stereotaxic surgery

Animals from five groups were anesthetized with intraperitoneal (i.p.) injection of Avertin solution (2,2,2-tribromoethanol, 240 mg/kg). For each animal, the bilateral intracranial injection of 0.6 μ g of AAV-VPS26B or vehicle (1.66 × 10^10^ vector genome/ μ l) into M1 on each hemisphere at 0.55 μL/min was performed using a stereotaxic apparatus. The coordinates from Bregma levels were anterior-posterior: +1.55 mm, medial-lateral: ± 1.65 mm, dorsal-ventral: 0.85 mm according to mouse brain atlas [27]. After injection, the syringe was maintained in position for 5 minutes to allow diffusion of the injected solution before being gently removed from the brain. Postoperatively, mice received saline (i.p.) to prevent dehydration. Mice were monitored regularly following recovery from surgery and were administered MPTP at 30 days following AAV-VPS26B or vehicle delivery.

### 2.3. MPTP administration

On the 30^th^ day of surgery, mice received i.p. injections of MPTP (20 mg/kg; Sigma-Aldrich, St. Louis, MO, USA) at two-hour intervals for a total of four injections.

### 2.4. Behavior test

The accelerating rotarod test was conducted on days 2, 3, and 5 following MPTP administration. Mice were placed on an automated rotarod apparatus and allowed to acclimate at 4 rpm for approximately 30 seconds. Animals then underwent three training trials (5 minutes each) during which the rotation speed was increased from 4 to 40 rpm. During testing, latency to fall was recorded with a cutoff time of 300 s. Each mouse performed three consecutive trials, and the average values were used for the analysis. All training and testing were conducted after MPTP injection to minimize the influence of pre-existing motor learning prior to dopamine depletion [28]. The control group (G1) served as the baseline.

### 2.5. Immunohistochemistry

The right cerebral hemispheres were sectioned into 30 μm free-floating coronal slices using a cryostat (Leica Biosystems) for histological analysis. Sections were treated to quench endogenous peroxidase activity in 3% hydrogen peroxide in methanol/PBS (1:1) for 15 minutes at RT, followed by incubation in methanol/PBS (1:1) for 10 minutes. After washing, sections were incubated in 4% normal horse serum containing 0.3% Triton X-100 for blocking. Brain sections were incubated with primary antibodies overnight at 4°C, followed by incubation with HRP-conjugated secondary antibodies for 2 hours at RT (Supplementary Table 2). For surface protein labeling, detergents were omitted from all solutions to prevent antibody penetration into cells [29]. Immunostaining was developed using DAB peroxidase substrate kit (Sera-care) according to the manufacturer’s instructions. Sections were mounted onto glass slides, dehydrated, and coverslipped.

For quantification of protein expression, images were obtained at 20× magnification with a digital light microscopy (PreciPoint M8, PreciPoint, Germany). The optical density (OD) of immunoreactivity in M1 and striatum was measured using ImageJ software. For quantification of TH-positive neurons, positive cells in the SN and VTA were counted manually using ViewPoint Light software. Results were presented as the number of cells per mm².

### 2.6. Immunofluorescence

Sections were blocked in 4% normal horse serum containing 0.3% Triton X-100 for 2 hours at RT, followed by incubation with primary antibodies overnight at 4 ° C. Sections were then incubated with secondary antibodies for 2 hours at RT (Supplementary Table 2). To label surface proteins in immunofluorescence, detergents were omitted from all solutions to prevent antibody penetration into cells [29]. Sections were mounted with mounting medium with DAPI (Vector Laboratories) for nuclear counterstaining.

### 2.7. Cell differentiation and viral infection

To induce dopaminergic differentiation, the human neuroblastoma cell line SH-SY5Y (ATCC, Manassas, VA) was sequentially treated with 10 μM retinoic acid (RA; Abcam) for 72 hours, followed by 80 nM 12-O-tetradecanoylphorbol-13-acetate (TPA; Abcam) for an additional 72 hours. During differentiation, cells were cultured in DMEM containing 2% heat-inactivated FBS, 100 U/ml penicillin, and 100 μ g/ml streptomycin under the same incubation conditions.

For overexpression of HA-tagged VPS26B, a total of 1 × 10⁶ SH-SY5Y cells were seeded in 60 mm dishes (Corning). After 24 hours of culture, cells were infected with AAV9-CAG-hVPS26B-HA (VectorBuilder) at a multiplicity of infection (MOI) of 10,000.

Mouse primary cortical neurons were kindly provided by Dr. Da Young Lee (Korea Research Institute of Bioscience and Biotechnology)

### 2.8. Co-Immunoprecipitation

Co-immunoprecipitation (Co-IP) was performed using the Pierce ™ Magnetic HA-Tag IP/Co-IP Kit (Thermo Fisher Scientific) according to the manufacturer’s instructions. Lysates from differentiated SH-SY5Y cells and mouse primary cortical neurons were incubated with anti-HA magnetic beads, and HA-tagged proteins were isolated using a magnetic stand (Thermo Fisher Scientific) and eluted following the kit protocol.

### 2.9. Western blot

Mouse brain tissues and cells were lysed in PRO-PREP ™ Protein Extraction Solution (iNtRON Biotechnology). After centrifugation, the supernatant was collected as the total protein lysate. Protein samples were mixed with 5X SDS-PAGE loading buffer containing 2% SDS and 5% β-mercaptoethanol and heated at 95°C for 10 minutes. Equal amounts of protein from each sample were loaded onto 10% or 12% (w/v) SDS-PAGE gels at 80 V for 2 hours. Proteins were then transferred onto nitrocellulose membranes (GE healthcare life sciences, pore size 0.45 µm) at 90V for 2 hours. Membranes were blocked in 5% skim milk (BD Difco) for 1 hour at RT and incubated with primary antibodies (Table 2) overnight at 4°C. After washing, membranes were incubated with secondary antibodies conjugated to HRP for 2 hours at RT. Immunoreactive proteins were detected using an enhanced chemiluminescence (ECL) detection system (Thermo Scientific). Densitometric analysis of western blots was performed using ImageJ software. Protein immunoreactivity was normalized to β-actin as a loading control.

### 2.10. Statistical analysis

Each mouse brain sample was assayed in at least two independent experiments. Statistical analyses were performed using GraphPad Prism 10. Data are displayed as mean ± SEM. An unpaired nonparametric test was used for comparison between two groups. For comparisons among multiple groups, data were analyzed using one-way ANOVA followed by Šídák’s multiple-comparisons test. Behavioral data were analyzed using a mixed-effects model (REML) with group and day as factors to account for repeated measurements and missing values. Post hoc comparisons were performed using Tukey’s multiple-comparisons test. Statistical significance was defined as follows: * p < 0.05, ** p < 0.01, *** p < 0.001 and **** p < 0.0001.

## 3. Results

### 3.1. AAV9-VPS26B-HA is predominantly expressed in neurons of the primary motor cortex

To assess VPS26B expression, exogenous VPS26B was identified by colocalization of HA and VPS26B immunoreactivity, which was detected in the M1 of VPS26B-overexpressing groups (G3, G5). Endogenous VPS26B was identified by VPS26B immunoreactivity alone and was observed in WT groups without VPS26B overexpression (G1, G2). Neither exogenous nor endogenous VPS26B was detected in the KO mouse group without VPS26B overexpression (G4) (Fig. 1A).

**Figure 1.**
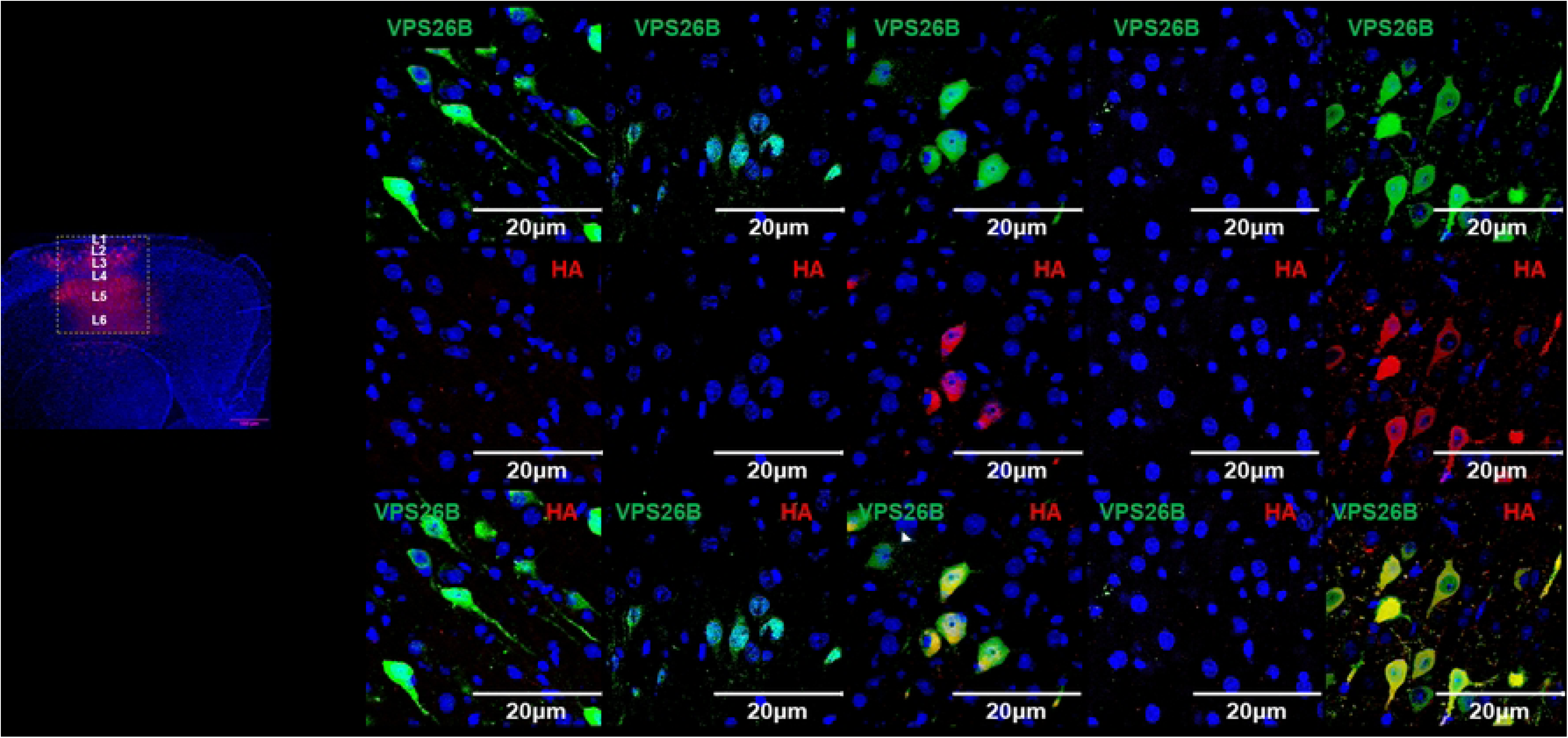
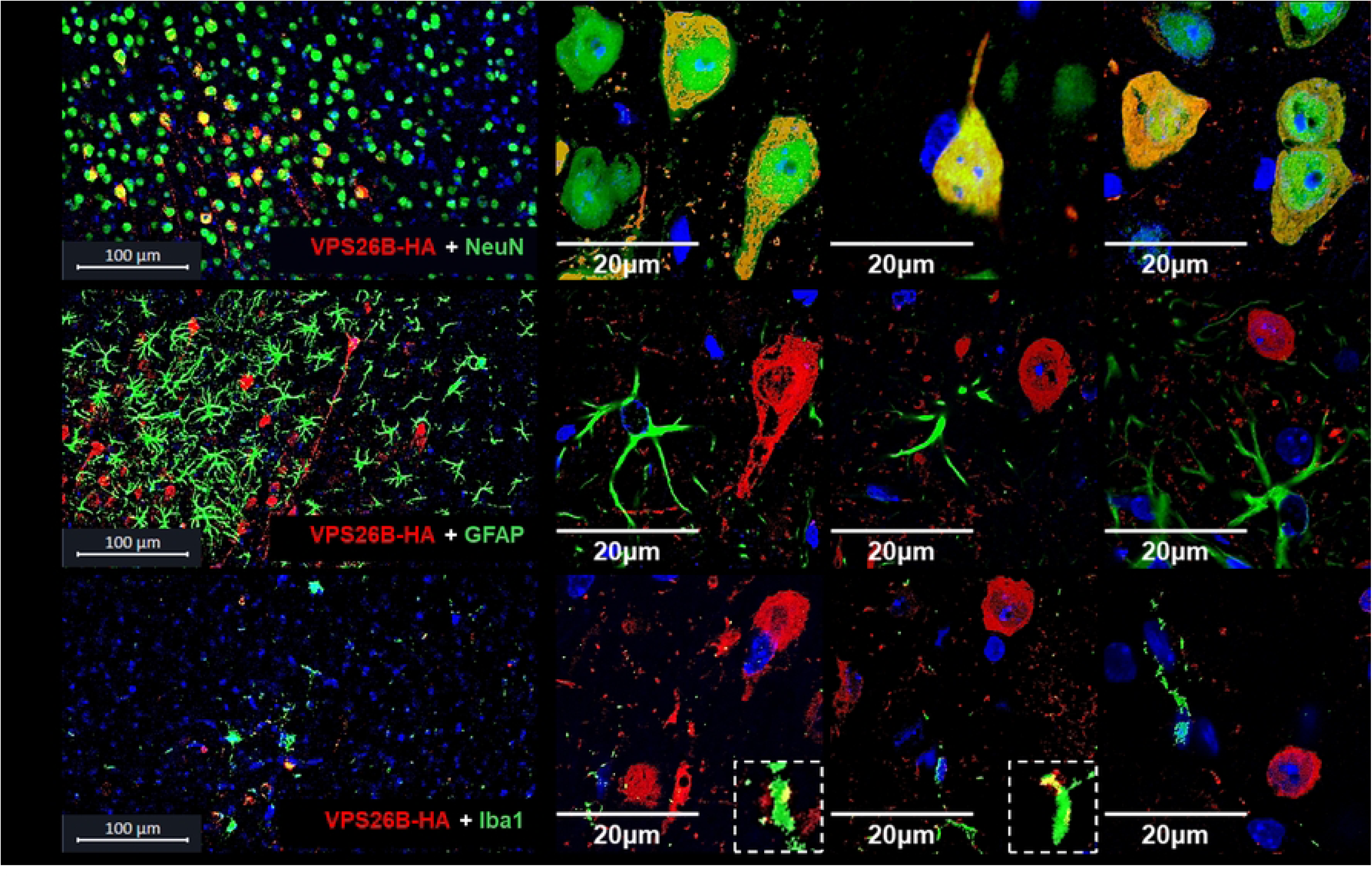
AAV-mediated VPS26B-HA is efficiently expressed in neurons of the primary motor cortex. **(A)** Representative immunofluorescence images showing endogenous VPS26B expression (VPS26B-positive cells only, green) and exogenous VPS26B expression (co-localization of VPS26B and HA tag; green and red, respectively) in the primary motor cortex (M1) across the five mouse groups. Scale bar= 100 µm and 20 µm. **(B** - **D)** Double immunofluorescence staining showing co-localization of HA (red) with cell-type markers (green) including NcuN (neuronal marker), GFAP (astrocyte marker) or Iba1 (mieroglial marker). Scale bar = 100 µm and 20 µm. Nuclei were counterstained with DAPI (blue). WT, wild type; KO, knockout; 26B O/E, VPS26B overexpression.

Co-immunofluorescent staining for HA with the cell-specific markers showed a strong colocalization of HA with neuronal marker NeuN (Fig. 1B), a negligible overlap with microglial marker Iba1, but not with astrocyte marker GFAP (Fig. 1C, D). Furthermore, a strong colocalization between HA and glutamatergic marker VGLUT2 indicated that exogenous VPS26B was delivered to glutamatergic neurons (Supplementary Figure 1).

### 3.2. Viral vector-mediated VPS26B overexpression partially preserves dopaminergic markers in MPTP-induced PD mouse model

To validate the Parkinsonian model, dopaminergic degeneration in the SN, VTA, striatum and M1 was assessed by tyrosine hydroxylase (TH) immunostaining. Compared with saline-treated controls (G1), MPTP-treated groups (G2-G5) exhibited a marked reduction of TH-positive cell number in the SN and VTA (Fig. 2A, B, D). Consistently, in the striatum and M1, TH-positive fiber density was significantly reduced in MPTP-treated groups (G2-G5) compared with controls (Fig. 2A, C, E). Notably, among MPTP-treated mice, TH-positive neuron numbers and fiber density were significantly higher in VPS26B-overexpressing groups (G3 and G5) than in non-overexpressing groups (G2 and G4) (Fig. 2A, F – I).

**Figure 2.**
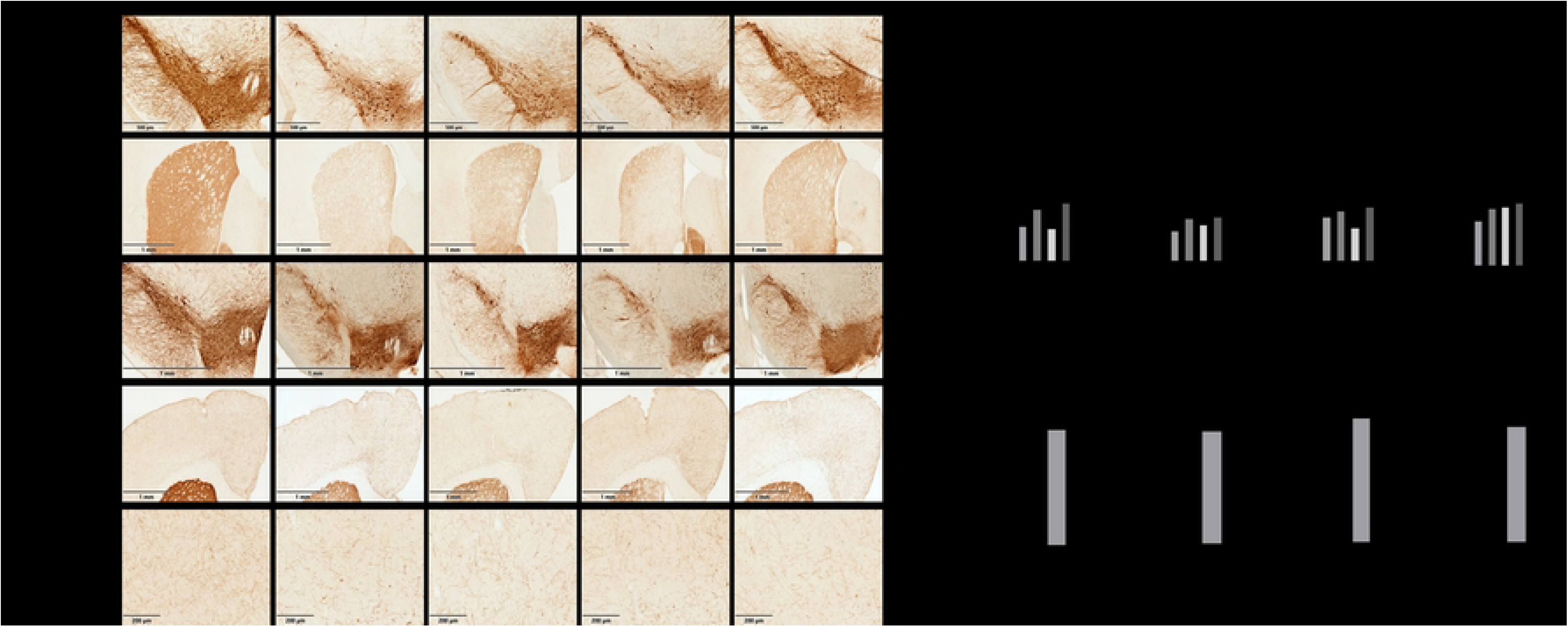
Viral vector-mediated VPS26B overexpression partially preserves dopaminergic markers in an MPTP-induced PD mouse model. **(A)** Representative immunohistochemistry images showing TH-positive dopaminergic neurons and fiber density in the SN, striatum, VTA and M1 in the five mouse groups. **(B, D)** Quantification of TH-positive dopaminergic neuron numbers in the SN and VTA across the five mouse groups. **(C, E)** Quantification of the optical density of TH immunoreactivity in the striatum and M1 across the five mouse groups. **(F, H)** Comparison of TH-positive neuron numbers in the SN and VTA between MPTP-treated groups with VPS26B overexpression and those without VPS26B overexpression. **(G, I)** Comparison of TH immunoreactivity optical density in the striatum and MI between MPTP-treated groups with VPS26B overexpression and those without VPS26B overexpression. Scale bar: SN, 500 µm; striatum, 1 mm; VTA, 1 mm; M1, 1 mm and 200 µm. Data are presented as mean ± SEM. Sample sizes: n = 4 (G1, G3, G5), n = 6 (G2), n = 3 (G4) mice per group. Statistical significance was determined using one-way ANOVA followed by Šidák’s multiple-comparisons test. ns, not significant, *p < 0.05, **p < 0.01, ***p < 0.001, and ****p < 0.0001. TH, tyrosine hydroxylase; SN, substantia nigra; VTA, ventral tegmental area; M1, primary motor cortex; WT, wild type; KO, knockout; 26B O/E, VPS26B overexpression.

These data confirm the dopaminergic degeneration following MPTP treatment and demonstrate that VPS26B overexpression partially attenuates dopaminergic neuronal and fiber loss in nigrostriatal as well as cortical regions.

### 3.3. MPTP reduces VPS26B in the primary motor cortex

One-way ANOVA revealed a significant effect of group on VPS26B levels (F(4, 37) = 63.31, p < 0.0001). Post hoc analysis showed that VPS26B levels were significantly reduced in WT mice following MPTP administration (G2 vs G1, p = 0.0103) and were absent in KO mice (G4). AAV-mediated VPS26B overexpression increased cortical VPS26B to 2.5-fold above baseline (Fig. 3A, B).

**Figure 3.**
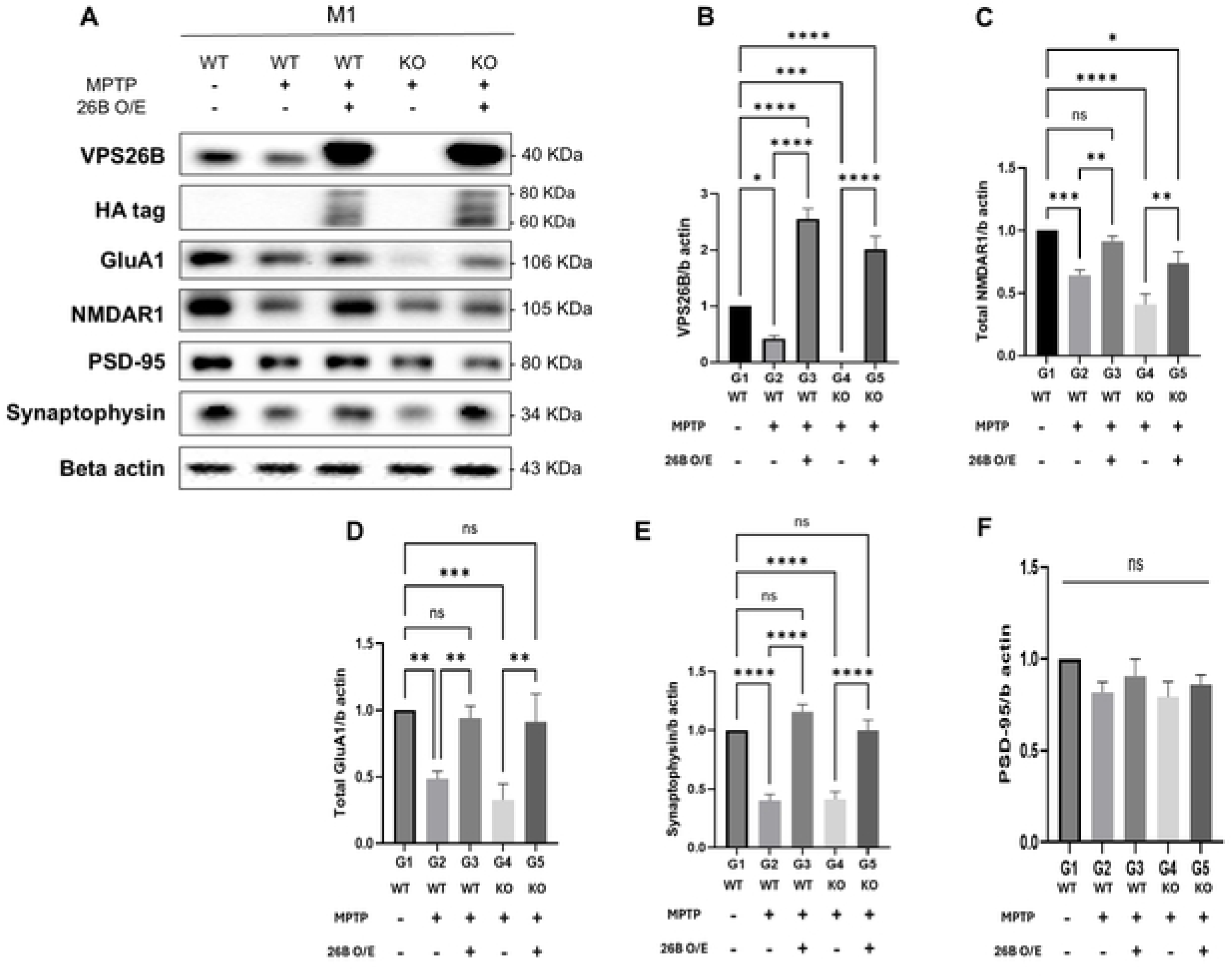
VPS26B overexpression attenuates MPTP-induced reductions in glutamate receptors and synaptic proteins in the primary motor cortex. **(A)** Representative western blot images showing VPS26B, NMDAR1, GluA1, PSD-95, and synaptophysin levels in the M1 of the five mouse groups. **(B - F)** Quantification of VPS26B, NMDAR1, GluA1, synaptophysin and PSD-95 levels in the M1 across the five mouse groups. Data are presented as mean ± SEM. Sample sizes: n = 4 (G1, G3, G5), n = 6 (G2), n = 3 (G4) mice per group. Statistical significance was determined using one-way ANOVA followed by Šidák’s multiple-comparisons test. ns, not significant, *p < 0.05, **p < 0.01, ***p < 0.001, and ****p < 0.0001. WT, wild type; KO, knockout; 26B O/E, VPS26B overexpression.

### 3.4. VPS26B overexpression attenuates MPTP-induced reductions in glutamate receptor and synaptic protein levels in the primary motor cortex

We examined whether changes in VPS26B expression correspond to alterations in glutamate receptor expression in the M1. One-way ANOVA revealed a significant effect of group on total NMDAR1 (F(4, 35) = 14.39, p < 0.0001) and GluA1 levels (F(4, 30) = 9.85, p < 0.0001). Post hoc analysis showed that MPTP treatment led to a significant decline in total NMDAR1 and GluA1 levels in WT mice compared with controls (G2 vs G1, p = 0.0089 and p = 0.0006, respectively), with a more pronounced reduction observed in KO mice (G4 vs G1, p < 0.0001). In contrast, these reductions were attenuated in VPS26B-overexpressing groups following MPTP administration (G3, G5) (Fig. 3A, C, D).

Surface GluA1 expression differed significantly among experimental groups (one-way ANOVA, F(4, 216) = 60.71, p < 0.0001). Post hoc analysis showed that surface GluA1 levels decreased significantly after MPTP injection in both WT and KO mice (G2 vs G1 and G4 vs G1, both p < 0.0001). In contrast, expression was preserved in VPS26B-overexpressing groups (G3 and G5), with no significant differences compared with controls. VPS26B overexpression significantly increased surface GluA1 levels relative to their respective MPTP-treated groups (G3 vs. G2 and G5 vs. G4, both p < 0.0001) (Fig. 4). These results indicate that VPS26B overexpression attenuates the MPTP-induced reduction in surface localization of GluA1 in M1.

**Figure 4.**
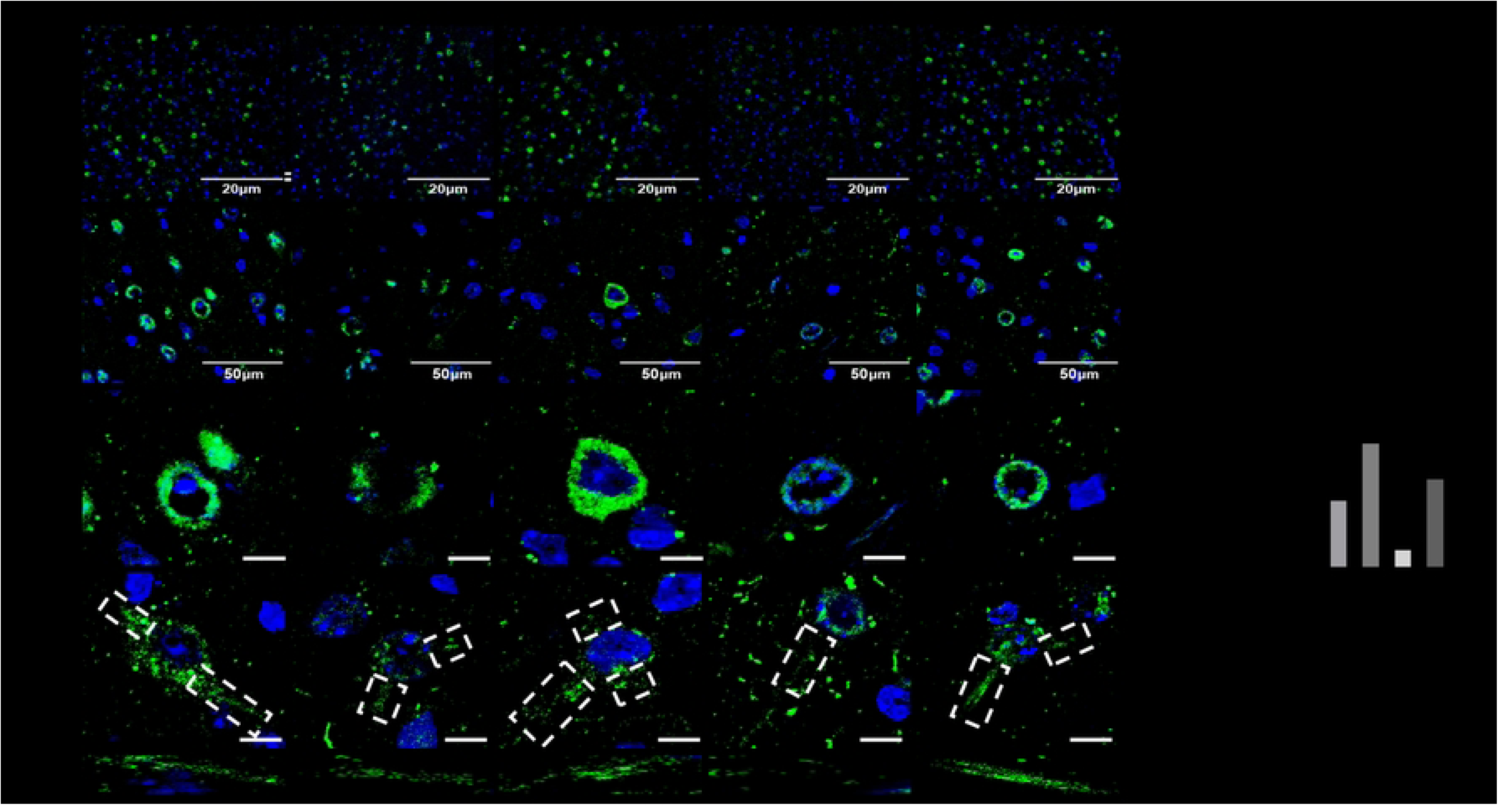
VPS26B overexpression attenuates MPTP-induced reduction 10 surface GluA1 expression in the primary motor cortex. **(A, B)** Representative immunofluorescence images showing surface GluA1 expression (green) in M1 across the five mouse groups. Scale bar= 50 µm (A) and 20 µm (B). **(C)** Representative image showing surface GluA1 localization around neuronal soma. **(D, E)** Representative images showing surface GluA1-positive puncta along neuronal processes (dashed boxes indicate representative regions) **(F)** Quantification of surface GluA1 fluorescence intensity per neuron in M1 across the five mouse groups Nuclei were counterstained with DAPI (blue). Data are presented as mean ± SEM. Sample sizes: n = 4 (G1, G3, G5), n = 6 (G2), n = 3 (G4) mice per group. Statistical significance was determined using one-way ANOVA followed by Šidák’s multiple-comparisons test. ns, not significant, *p < 0.05, **p < 0.01, ***p <0.001, and ****p <0.0001. M1, primary motor cortex; WT, wild type; KO, knockout; 26B O/E, VPS26B overexpression.

### 3.5. VPS26B overexpression mitigates MPTP-induced loss of synaptic proteins in the primary motor cortex

We next assessed the expression of synaptic plasticity-related proteins, including synaptophysin and PSD-95. One-way ANOVA revealed a significant effect of group on synaptophysin levels (F(4, 37) = 36.76, p < 0.0001). Post hoc analysis showed that cortical synaptophysin levels were significantly reduced following MPTP treatment in both WT and KO mice (G2 vs G1 and G4 vs G1, both p < 0.0001). In contrast, synaptophysin levels were maintained in groups receiving AAV-VPS26B overexpression (G3, G5) (Fig. 3A, E). No significant differences in PSD-95 levels were observed among groups (F(4, 37) = 1.57, p = 0.2015) (Fig. 3A, F). These results indicate that MPTP disrupts the expression of synaptic function-related proteins in the M1, with a more pronounced effect on the presynaptic marker synaptophysin, and that VPS26B overexpression mitigates the synaptic protein loss caused by MPTP.

### 3.6. VPS26B is associated with altered motor performance under Parkinsonian conditions

We next examined motor learning using the accelerating rotarod test. Mixed-effects analysis (REML) revealed significant main effects of day (F(1.74, 106.2) = 19.84, p < 0.0001) and group (F(4, 64) = 15.01, p < 0.0001), as well as a significant day × group interaction (F(6.97, 106.2) = 4.23, p = 0.0004), indicating that motor learning trajectories differed across groups.

Following MPTP administration, WT mice (G2) exhibited reduced performance at early time points compared with controls (G2 vs G1, p < 0.0001), followed by gradual improvement across testing sessions. VPS26B overexpression in WT mice (G3) showed similar improvement, with a slight numerical increase that did not reach statistical significance (G3 vs G2, p = 0.8812). In contrast, VPS26B-deficient mice (G4) displayed variable performance across testing sessions, with a transient improvement between days 2 and 3, followed by a decline at day 5. Notably, VPS26B-overexpressing KO mice (G5) maintained performance at levels comparable to controls across testing sessions, although a trend toward lower performance was observed on the first day of testing but did not reach statistical significance (G5 vs G1, p = 0.4098) (Fig. 5). Statistical comparisons among the five groups are presented in Supplementary Figure 2.

**Figure 5.**
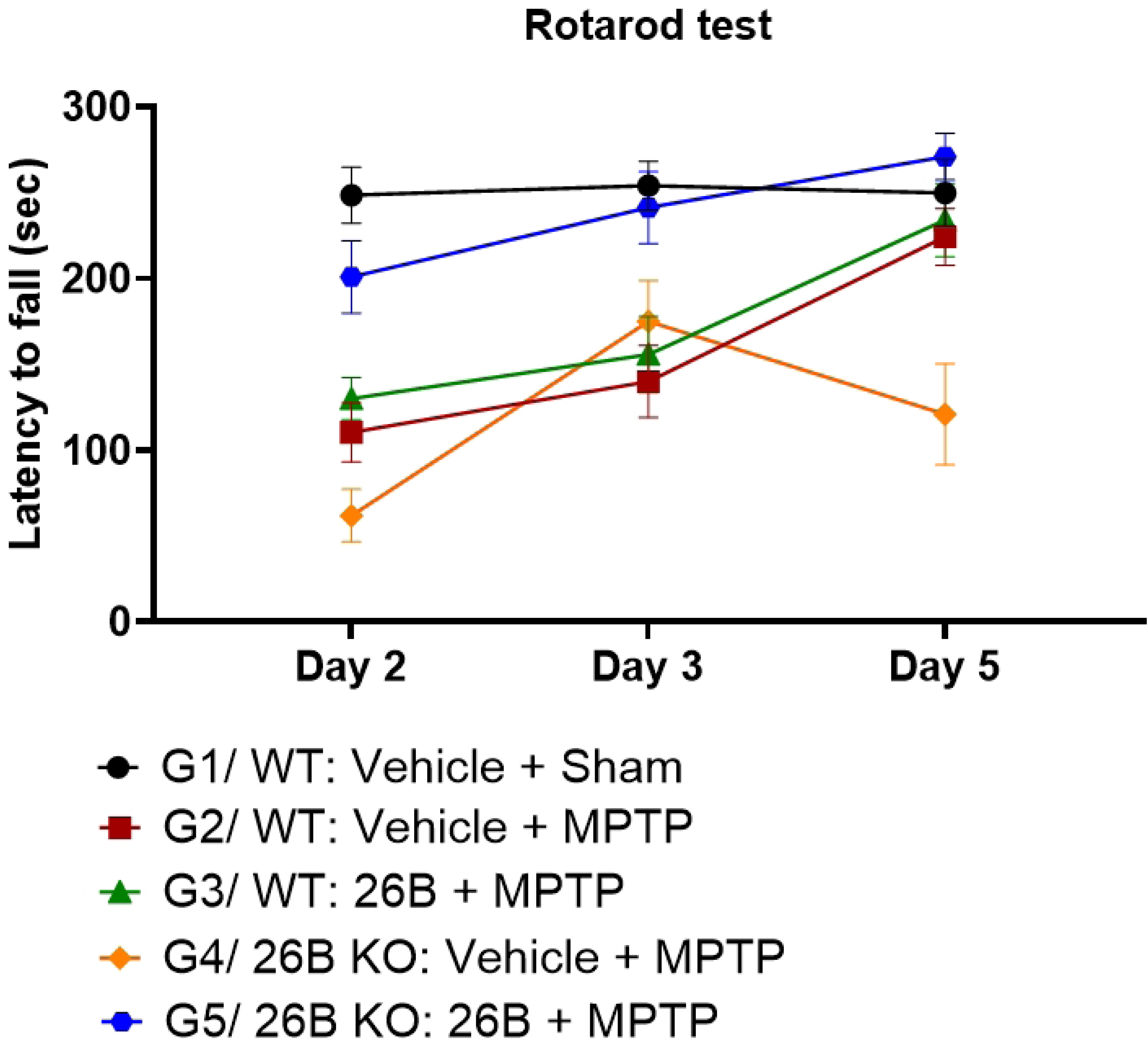
VPS26B deficiency alters rotarod performance under MPTP-induced conditions. Rotarod performance showing latency to fall during the accelerating rotarod test across the five mouse groups on days 2, 3, and 5 following MPTP administration. Data are presented as mean ± SEM. Behavioral data were analyzed using a mixed-effects model (REML) with group and day as factors, followed by Tukey’s multiple-comparisons test. Sample sizes: n = 4 (G1, G3, G5), n = 6 (G2), and n = 3 (G4) mice per group. Each mouse performed three trials per day, and the average latency was used for analysis.

### 3.7. VPS26B interacts with dopamine D1 and D2 receptors in SH-SY5Y dopaminergic cells, but not in primary cortical cells

To explore the cause of VPS26B reduction under dopamine deficiency, we performed co-IP followed by western blot to assess the interaction between VPS26B and dopamine receptors D1 and D2. HA-tagged and VPS35 were used as positive controls. Co-immunoprecipitation revealed that VPS26B interacts with both dopamine D1 and D2 receptors in SH-SY5Y cells, whereas no interaction was detected in primary cortical neurons, suggesting a potentially cell type–dependent interaction (Fig. 6). Furthermore, we attempted to detect the interaction between VPS26B and GluA1; however, no association was observed in either dopaminergic cells or primary cortical cells (data not shown).

**Figure 6.**
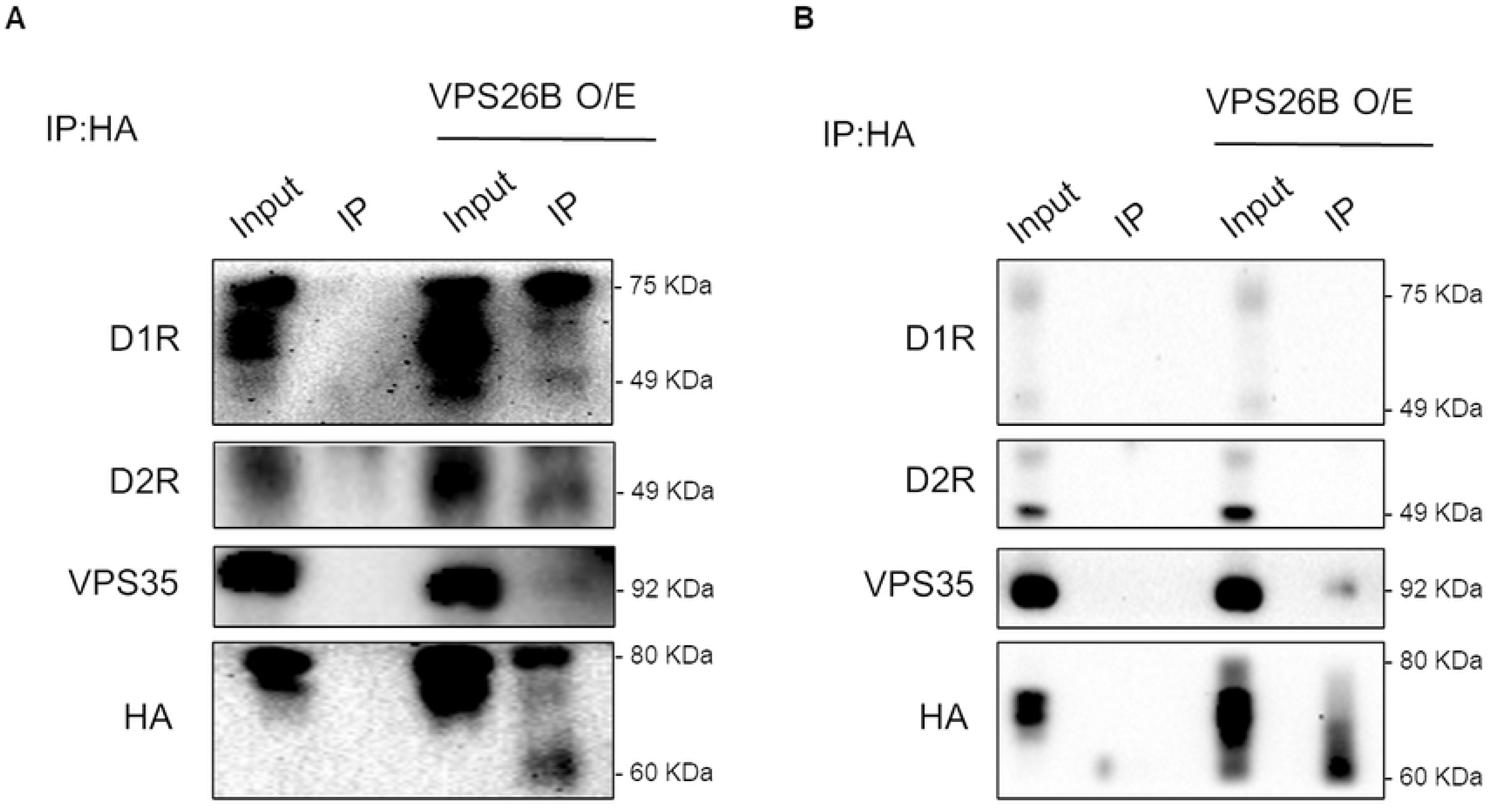
VPS26B interacts with dopamine D1 and D2 receptors in differentiated SH-SY5Y cells, but not in primary cortical neurons. Co-immunoprecipitation of HA-tagged VPS26B from lysates of **(A)** differentiated SH-SYSY cells and **(B)** primary cortical neurons, followed by western blot analysis for D1R, D2R, VPS35, and HA. VPS35 and HA were used as positive controls. Blots shown are representative of at least two independent experiments. IP, immunopreeipitation; O/E, overcxprcssion.

## 4. Discussion

In this study, we identify VPS26B as a potential modulator of cortical GluA1 and synaptic protein expression. Following MPTP administration, VPS26B deficiency in the mouse M1 coincided with reduced surface GluA1 expression, synaptic protein loss, and altered motor performance, whereas VPS26B overexpression attenuated these alterations.

Our data support a model in which cortical VPS26B deficiency impairs maintenance of surface GluA1 levels. These findings are consistent with previous evidence demonstrating endosomal trapping of GluA1 in the trans-entorhinal cortex region under conditions of VPS26B depletion [24]. Under these conditions, cargo proteins may be preferentially directed toward lysosomal degradation [30, 31], which aligns with the reduction in total GluA1 in our study. These reductions were prevented by VPS26B overexpression in M1. In addition to its role in synaptic signaling, GluA1 has been implicated in neurite outgrowth and structural plasticity, raising the possibility that VPS26B may influence cortical circuitry through additional GluA1-related mechanisms [32–34].

MPTP also reduced cortical NMDAR1 levels, with a more pronounced decrease in VPS26B-deficient mice. Given that NMDA receptor activity has been implicated in the regulation of GluA1 insertion via calcium influx [35], the reduction in NMDAR1 may further contribute to the decreased surface expression of GluA1. Consistent with this, NMDAR1 in M1 is known to play a role in activity-dependent synaptic strengthening and associative learning [8]. VPS26B overexpression attenuated the reduction in NMDAR1 levels, suggesting that VPS26B may contribute to maintaining glutamate receptor expression in the MPTP model.

Surface GluA1 expression has been linked to increased presynaptic synaptophysin via a retrograde signal, coordinating the formation or strengthening of nascent synapses [36, 37]. Synaptophysin is a key presynaptic protein required for synaptic function, and its loss impairs synaptic plasticity, as shown in synaptophysin knockout mice [38]. Reduced synaptophysin levels have also been reported in PD models and patients [39–41]. In our MPTP model, VPS26B overexpression preserved synaptophysin levels, possibly through preservation of surface GluA1 availability. This preservation may contribute to the improved rotarod performance in MPTP-treated mice.

During behavioral testing, MPTP-treated mice exhibited reduced rotarod performance at early time points compared with controls. VPS26B overexpression mitigated these deficits, whereas VPS26B deficiency exacerbated performance instability across testing sessions. In addition to assessing motor coordination, the accelerating rotarod test also provided insights into motor learning through changes in performance over testing sessions [42]. In the present study, behavioral testing was conducted after MPTP administration without prior training, minimizing recall and use of stored motor information prior to dopamine depletion [28]. Taken together, VPS26B overexpression coincided with improved rotarod performance under MPTP conditions.

VPS26B overexpression also showed a trend toward attenuating dopaminergic degeneration. These findings are consistent with previous reports suggesting that cortical alterations may influence dopaminergic vulnerability [4]. In this context, preservation of cortical synaptic proteins by VPS26B may indirectly support dopaminergic neuron survival. Additionally, co-immunoprecipitation experiments suggested a possible interaction between VPS26B and D1R/D2R in neuronal models. Although the functional significance of this interaction remains to be established, these observations suggest a potential link between VPS26B and dopaminergic signaling under dopamine-depleted conditions.

Several limitations should be noted. First, VPS26B knockout control group under saline treatment was not included, which may limit the interpretation of baseline effects of VPS26B deficiency. Second, synaptic function was assessed through the expression of synaptic plasticity–related proteins. Future research should involve electrophysiological recordings to directly verify alterations in cortical synaptic plasticity. Finally, the relatively small sample size in several experimental groups, particularly following MPTP-associated mortality, may limit statistical power.

In summary, impairments in surface glutamate receptor and synaptic protein expression in the MPTP model are at least partially attributable to a loss of VPS26B in M1, which may contribute to motor learning deficits. VPS26B overexpression in the M1 attenuated these impairments, suggesting VPS26B may represent a candidate target for modulating synaptic dysfunction and motor learning impairment in PD.

